# Short-term plasticity persists in the absence of PKC phosphorylation of Munc18-1

**DOI:** 10.1101/2021.02.14.431144

**Authors:** Chih-Chieh Wang, Christopher Weyrer, Diasynou Fioravante, Pascal S. Kaeser, Wade G. Regehr

## Abstract

Post tetanic potentiation (PTP) is a form of short-term plasticity that lasts for tens of seconds following a burst of presynaptic activity. It has been proposed that PTP arises from protein kinase C (PKC) phosphorylation of Munc18-1, an SM (Sec1/Munc-18 like) family protein that is essential for release. To test this model, we made a knockin mouse in which all Munc18-1 PKC phosphorylation sites were eliminated through serine-to-alanine point mutations (Munc18-1 SA mice). Expression of Munc18-1 was not altered in Munc18-1SA mice, and there were no obvious behavioral phenotypes. At the hippocampal CA3 to CA1 synapse, and the granule cell parallel fiber to Purkinje cell (PF to PC) synapse, basal transmission was largely normal except for small decreases in paired-pulse facilitation that are consistent with a slight elevation in release probability. Phorbol esters that mimic activation of PKC by diacylglycerol still increased synaptic transmission in Munc18-1 SA mice. In Munc18-1 SA mice, 70% of PTP remained at CA3 to CA1 synapses, and the amplitude of PTP was not reduced at PF to PC synapses. These findings indicate that at both CA3 to CA1 and PF to PC synapses, phorbol esters and PTP enhance synaptic transmission primarily by mechanisms that are independent of PKC phosphorylation of Munc18-1.

**Significance Statement:** A leading mechanism for a prevalent form of short-term plasticity, post-tetanic potentiation (PTP), involves protein kinase C phosphorylation of Munc18-1. This study tests this mechanism by creating a knock in mouse in which Munc18-1 is replaced by a mutated form of Munc18-1 that cannot be phosphorylated. The main finding is that most PTP at hippocampal CA3 to CA1 synapses, or at cerebellar granule cell to Purkinje cell synapses does not rely on PKC phosphorylation of Munc18-1. Thus, mechanisms independent of PKC phosphorylation of Munc18-1 are important mediators of PTP.

## Introduction

A brief burst of presynaptic activity transiently increases synaptic strength for tens of seconds at many types of synapses (Zucker and Regehr, 2002). In some cases, the resulting enhancement is divided into a short-lived component (augmentation) and a longer-lived component (post-tetanic potentiation, PTP). Because distinguishing between augmentation and PTP is difficult, we refer to all transient post-tetanic synaptic enhancement lasting for seconds to tens of seconds as PTP. It has been proposed that PTP contributes to short-term memory (Vandael et al., 2020; Vyleta et al., 2016; Zucker and Regehr, 2002)

Clarifying the mechanism of PTP is important in its own right, and could lead to a means of selectively eliminating PTP to provide insight into its functional roles. One of the most promising mechanisms for PTP is that calcium activates protein kinase C (PKC) that in turn phosphorylates Munc18-1 to increase neurotransmitter release (Brager et al., 2003; Fioravante et al., 2011; Genç et al., 2014; Wierda et al., 2007). Munc18-1 plays an essential role in neurotransmitter release. The current working model is that Munc18-1 initially binds the close conformation of syntaxin-1, and then along with Munc13-1 opens syntaxin-1 leading to partial SNARE complex assembly and the formation of the readily releasable pool (Arunachalam et al., 2007; Dawidowski and David, 2013, 2016; Dulubova et al., 1999; Imig et al., 2014; Rizo, 2018; Toonen et al., 2005). Deletion of Munc18-1 eliminates the formation of the RRP and eliminates both spontaneous and evoked neurotransmitter release (Aravamudan et al., 1999; Richmond et al., 1999; Varoqueaux et al., 2002; Verhage et al., 2000). It has been proposed that phosphorylation of Munc18-1 regulates neurotransmitter release (Craig et al., 2003; de Vries et al., 2000; Genç et al., 2014; Wierda et al., 2007). The initial evidence in favor of PKC involvement in PTP was that PKC inhibitors eliminate PTP, and activators (phorbol esters) occlude PTP (Alle et al., 2001; Beierlein et al., 2007; Brager et al., 2003; Korogod et al., 2007; Lee et al., 2007; Rhee et al., 2002; Wierda et al., 2007). Calcium-dependent PKC isoforms have been implicated in PTP at the calyx of Held (Fioravante et al., 2011) and the cerebellar PF to PC synapse (Fioravante et al., 2012). The replacement of Munc18-1 with a mutated version that cannot be phosphorylated by PKC eliminated PTP at cultured hippocampal synapses (Wierda et al., 2007) and reduced PTP at the calyx of Held (Genç et al., 2014). This manipulation also eliminated or strongly attenuated synaptic enhancement produced by phorbol esters (Genç et al., 2014; Wierda et al., 2007). Previous studies also implicated a number of other proteins in PTP, including calmodulin, CaM kinase II, myosin light chain kinase, Munc13, Synaptotagmin 1, and several other candidates (Chapman et al., 1995; De Jong et al., 2016; Fiumara et al., 2007; Junge et al., 2004; Lee et al., 2008; Rosahl et al., 1995; Vandael et al., 2020; Wang and Maler, 1998).

Our goal was to test the hypothesis that PKC phosphorylation of Munc18-1 is essential to a general mechanism of PTP, and we focused on hippocampal CA3 to CA1 synapses and cerebellar PF to PC synapses. Both of these synapses exhibit PTP that is thought to involve PKC (Brager et al., 2003; Fioravante et al., 2012). We could not use the approaches that had been taken to study cultured hippocampal neurons in which mutated Munc18-1 was expressed in neurons from Munc18-1 knockout animals (Wierda et al., 2007), because Munc18-1 knockout mice are embryonically lethal and individual neurons lacking Munc18-1 die (Verhage et al., 2000). In addition, CA1 pyramidal cells and cerebellar Purkinje cells receive inputs from a great many presynaptic cells, and it is difficult to express rescue constructs in a large fraction of presynaptic cells. We therefore generated a knockin mouse (Munc18-1SA mouse) in which Munc18-1 is replaced with a mutated form that cannot be phosphorylated by PKC in order to study the contribution of PKC phosphorylation of Munc18-1 to PTP. We find that in Munc18-1SA mice, PTP and enhancement by phorbol esters are still largely intact for both CA3 to CA1 and PF to PC synapses. This suggests that PKC phosphorylation of Munc18-1 is not the primary mechanism underlying PTP at CA3 to CA1 and PF to PC synapses.

## Material and Methods

### Animals

All animal experiments were completed in accordance with guidelines set by the Harvard Medical Area Standing Committee on Animals. PKCαβγ triple knockout (TKO) mice were obtained through breeding of PKCα, PKCβ, and PKCγ knockout animals (Abeliovich et al., 1993; Leitges et al., 2002; Leitges et al., 1996).

### Generation of Munc18-1SA mice

Serines 306, 312 and 313, which are in exon 11 of the Munc18-1 gene were replaced with non-phosphorylatable alanines using homologous recombination at InGenious Targeting Laboratory (Long Island, NY). Briefly, a targeting vector was constructed from a 9.53 kb fragment of a C57BL/6 BAC clone (RP23: 320N1). The construct contained from 5’ to 3’: a 6.7 kb long homology arm, a conditional inversion cassette with a wild-type exon 11, a neomycin resistance gene and an inverted SA exon 11 flanked by loxp/lox2272 sites, a short homology arm, and a diphtheria toxin expressing cassette. The targeting vector was confirmed by restriction analysis and sequencing. Hybrid (129/Sv × C57BL/6) embryonic stem cells were electroporated with the NotI-linearized targeting vector, and homologous recombined ES cell clones were identified by PCR and sequencing. Upon expansion, homologous recombination was confirmed by Southern blotting using BclI digestion of genomic DNA and a 5’ outside probe. Chimeric mice were generated by blastocyst injections and bred to C57BL/6 mice. Following confirmation of germline transmission of the mutant allele, the neomycin resistance cassette was removed by crossing the mice with flp recombinase transgenic mice (Dymecki, 1996) to generate the Munc18-1cSA allele. Constitutive knockin mice (Munc18-1SA mice) were generated by Cre recombination in the germ line using human β − actin-Cre mice. The Munc18-1SA mice were genotyped by PCR with oligonucleotide primers (5’-TTG GAG TAG GAA TAC TGG CCC −3’ and 5’-ACA GAA GAG GAG CTG ACC CCT G −3’ to identify a 573 bp KI band and a 511 bp wild-type band. DNA sequencing (Eton Biosciences) and protein mass spectrometry analysis (Harvard Mass Spectrometry and Proteomics Resource Laboratory) confirmed that wild-type Munc18-1 had been replaced by Munc18-1 containing point mutations that eliminate PKC phosphorylation sites. Experiments were performed using Munc18-1SA mice and animals that express wildtype Munc18-1.

### Measurement of motor activity

Mice between 3 and 4 weeks of age were used for motor activity assays. All behavioral experiments were conducted in the dark phase of the light-dark cycle. Individual mice were placed in a plastic chamber (23 × 16 inches) filled with standard mouse bedding. Open field activity was recorded using a USB IR-illumination camera (ELP) in near-darkness for 10 minutes shortly after being placed in the chamber. The chamber was thoroughly cleaned with ethanol after each trial and the bedding was replaced. Mouse behavior was tracked using custom scripts written in Matlab. Analysis was conducted blind to genotype.

### Preparation of brain slices

Mice of either sex aged p12-15 days (for cerebellar slices) or p18-25 days (for hippocampal slices) were anesthetized with isoflurane and decapitated. Acute transverse slices (300-320 μm thick for hippocampal slices and 250 μm thick for cerebellar slices) were cut in ice-cold solution consisting of the following (in mM): 110 Choline-Cl, 7 MgSO_4_, 2.5 KCl, 1.2 NaH_2_PO_4_, 0.5 CaCl_2_, 25 glucose, 11.6 Na-Ascorbate, 2.4 Na-Pyruvate, and 25 NaHCO_3_. Slices were then incubated at 32°C for 20-30 min in a bicarbonate-buffered solution composed of the following (in mM): 125 NaCl, 25 NaHCO_3_, 1.25 NaH_2_PO_4_, 2.5 KCl, 1.5 CaCl_2_ (2 CaCl_2_ for cerebellar slices), 1.5 MgCl_2_ (1 MgCl_2_ for cerebellar slices) and 25 glucose. The slices were kept in the chamber at room temperature until recording or protein extraction.

### Electrophysiology

The recordings for hippocampal CA3 to CA1 synapses were performed as described previously (Wang et al., 2016). Briefly, recordings were conducted at 30-32 °C, and the hippocampal CA3 region was cut from the CA1 region with a scalpel blade to prevent recurrent excitation. The external solution was the same solution used for incubating slices but supplemented with the following drugs (in μM): 20 bicuculline, 2 CGP, 2 CPP, and 1 AM251. A stimulation electrode filled with ACSF was placed at least 500 μm away from the recording site to stimulate the Schaffer collateral fibers. For field recordings, the recording pipettes (0.3-2 MΩ) were filled with ACSF and placed in the *stratum radiatum* in the CA1 dendritic area, and input-output curve with the stimulation range 10-80 μA was first analyzed to determine the stimulation intensity that roughly produced half of the maximum response without evoking population spikes. Recordings of granule cell parallel fiber to PC synapses were made similar as described previously (Fioravante et al., 2012). The internal consisted of 100 CsCl, 35 CsFl, 10 EGTA, 10 HEPES and 3 or 5 QX-314-Cl (Fioravante et al., 2012; Weyrer et al., 2019). PF to PC experiments was performed at 30-36°C.

mEPSCs were recorded in whole-cell configuration using 2 MΩ pipettes filled with the same internal solution used to measure evoked EPSCs. TTX (1 µM) was additionally added to the external solution to block action potential evoked synaptic currents. Data were recorded in 30 sec epochs, sampled at 2.5 kHz, and filtered at 500 Hz with an eight-pole Bessel filter (Frequency Devices, Haverhill, MA). Membrane potential was maintained at −70 mV. Series resistance compensation was not used. Leak currents were −10 to −200 pA. Events were counted and analyzed off-line using IGOR PRO software (Wavemetrics, Lake Oswego, OR) and custom macros provided by Monica Thanawala. Inclusion criteria were a 4–10 pA amplitude threshold, a minimum rate of rise of 0.4 pA/msec, and a decay time constant between 3 and 12 msec. At least 2 trials were recorded for each cell to obtain the average mEPSC frequency and amplitude.

For cerebellar experiments, stimuli were given every 2.5 s, then PTP-inducing 10 stimuli at 50 Hz were applied 2.5 seconds after the last EPSC of the baseline. One second after the train, stimuli were resumed and delivered every 2.5 s.

Experiments in Figure 5a and 5b were performed blind to genotype. Other data sets consist of blinded and unblinded experiments and are therefore considered unblinded to genotype. Data are expressed as mean ± SEM.

### Western-Blotting

Western-Blotting was conducted as described previously (Wang et al., 2016). Brains excluding cerebellum from animals of age at 2-3 weeks were extracted acutely and homologized in ice-cold lysis buffer containing 150mM NaCl, 25mM HEPES, 4 mM EGTA, phosphatase inhibitor (Roche #04906845001), and protease inhibitor (Sigma #P8340). Total protein concentration was determined by a BCA assay (Pierce), and 30 μg of protein in Laemmli sample buffer was loaded onto 10% polyacrylamide gels (Bio-Rad). After SDS-PAGE, gels were transferred onto nitrocellulose membranes and blocked in 5% nonfat milk (Cell Signaling Technology) in TBS-Tween 20 (TBST). Membranes were incubated overnight at 4°C with primary
antibodies, followed by the incubation with fluorescent secondary antibodies for 1.5 hour. Membranes were washed with TBST several times followed with two times wash in TBS. Membranes were dried and imaged in Odyssey Classic (LI-COR). Non-saturated images were quantified in ImageJ. Analysis was conducted blind to genotype. For analysis, the fluorescent intensities of target bands were measured blind to genotype and then normalized to β-Actin fluorescent intensity. The normalized value was further normalized to the averaged wildtype control values in the same gel. Data are expressed as mean ± SEM and statistical analysis were conducted using paired t tests. The following antibodies were used: Munc18-1 (1:1000; Abcam, #ab3451), β-Actin (1:5000, Sigma, #A1978), rabbit phospho- and fluorescent antibodies (1:10,000, IRDye 800CW Donkey anti-Rabbit IgG, LI-COR #926-32213 and IRDye 680RD Donkey anti-Mouse IgG, LI-COR #926-68072).

### Mass Spectrometry

2.5 μ l Munc18-1 antibody was added to 500 μg brain lysate, followed by the incubation at 4°C for overnight. Protein lysate was extracted in ice-cold lysis buffer containing 150 mm NaCl, 25 mm HEPES, 4 mm EGTA, phosphatase inhibitor (Roche, 04906845001), and protease inhibitor (Sigma-Aldrich, P8340). Total protein concentration was determined by a BCA assay (Pierce) 60 μl Protein A beads with 50% slurry (Cell Signaling Technology, #9863S) were then added and incubated for 6 hours to pull down the antibody-protein complex. Beads were then washed with PBS and subjected to SDS-PAGE. The gel was stained with Coomassie blue and the band that corresponds to Munc18-1 molecular weight was cut. That band was subjected to Mass Spectrometry analysis by Harvard Mass Spectrometry and Proteomics Resource Laboratory.

## Results

### Generation and initial characterization of Munc18-1SA mice

To study the role of PKC phosphorylation of Munc18-1, we used homologous recombination to generate conditional knockin mice in which Munc18-1 cannot be phosphorylated by PKC. This was achieved by using homologous recombination in embryonic stem cells to replace serines 306, 312 and 313 that can be phosphorylated by PKC (Craig et al., 2003; de Vries et al., 2000; Wierda et al., 2007), with non-phosphorylatable alanines (Munc18-1SA: S306A/S312A/S313A) (**Fig. 1A**). Successful recombination was confirmed first by PCR and DNA sequencing, and by Southern blotting (**Fig. 1B**). Chimeric mice with the Munc18-1 recombined (R) allele were obtained by blastocyst injection. Upon germline transmission, the neomycin-resistant cassette was removed by flp recombination using transgenic mice (Dymecki, 1996) to produce the conditional SA knockin allele (cSA) (**Fig. 1C**). Munc18-1cSA mice were hypomorphic, expressed only 23% of Munc18-1 in control mice (n=11, p<0.001) (**Fig. 1E**), and had 18% reduced body weight (n=21, p<0.001) (**Fig. 1F**). Cre-recombination was performed using transgenic mice (Lewandoski and Martin, 1997) to generate the constitutive Munc18-1 SA allele (SA), which recovered Munc18-1 expression levels and body weight to control, and the offspring of heterozygote matings survived at Mendelian ratios (of 258 mice 61 were WT, 69 were SA and 128 were heterozygotes). Sequencing of the Munc18-1 PCR product and mass spectrometry of brain protein extracts confirmed that serines 306, 312 and 313 were replaced by alanines in these mice (**Fig. 1F**).

**Figure 1.**
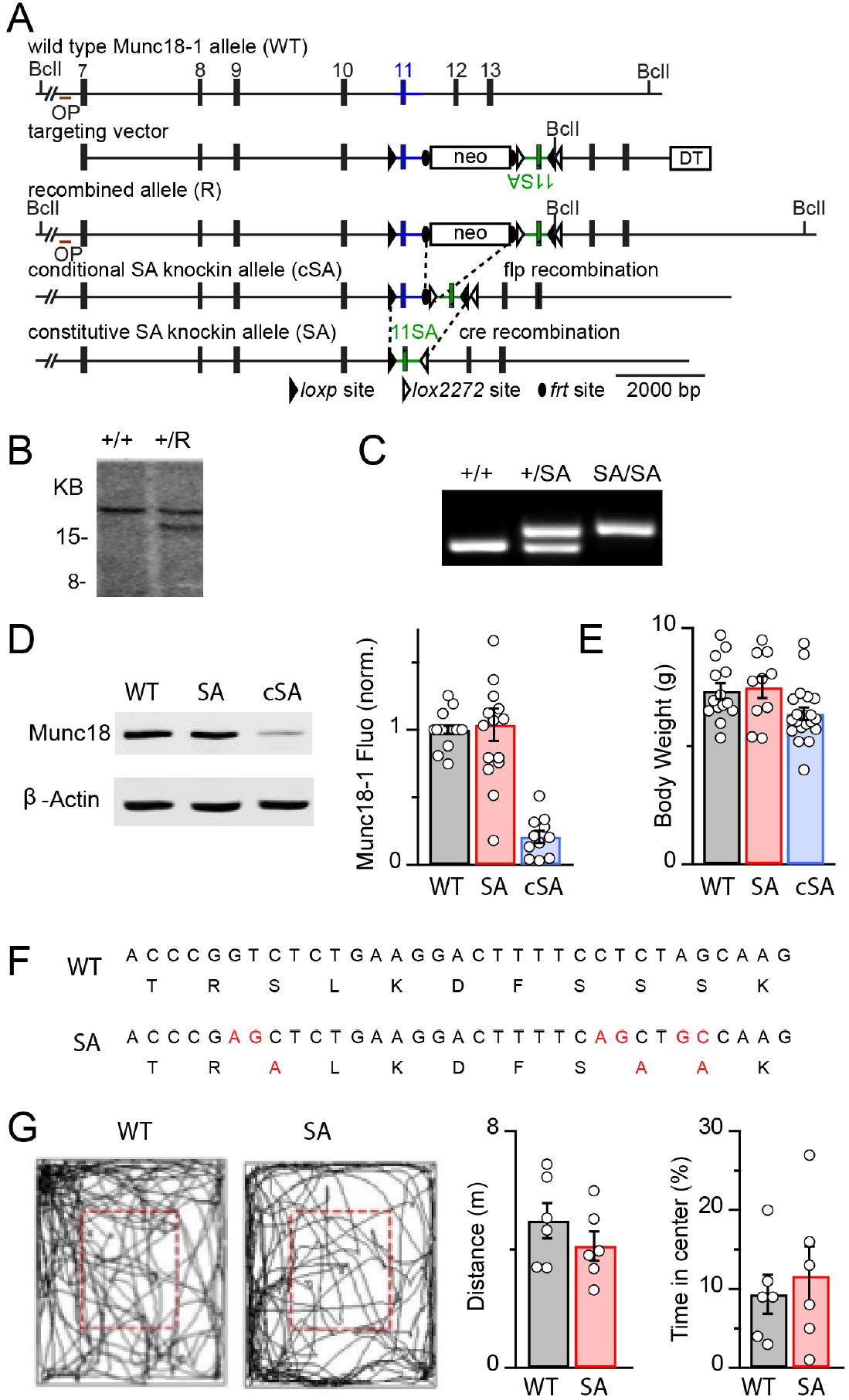
Generation and Basic Characterization of Munc18-1 SA mice. A. Knockin strategy for Munc18-1 SA-KI mice. Abbreviations: DT, diptheriatoxin-expressing cassette; neo, neomycin resistance cassette. B. Southern blot of DNA digested with BclII from embryonic stem cells from wildtype controls and heterozygous cells after homologous recombination. C. PCR genotyping of wildtype mice, heterozygous knockin SA mice, and homozygous knockin SA mice. D. DNA (top) and protein (bottom) sequencing are shown for wildtype and SA knockin mice with altered regions highlighted (red). E. Body weights of (P12-P15) wildtype, Munc18-1 SA and Munc18 cSA mice. F. (left) Western blots of brain homogenates for wildtype, Munc18-1 SA and Munc18-1 cSA mice. (right) Summary of Munc18-1expression levels assessed by western blotting with fluorescent secondary antibodies. G. (left) Open field assay examples for a wildtype and a Munc18-1 SA mouse. (right). Summaries of the total distance travelled in 10 minutes and the percentage of time spent in the center (within the red dashed line).

There were no obvious behavioral deficits. Motor activity was assessed in open field tests and was similar in littermate controls and Munc18-1SA mice (**Fig. 1G**). The distance traveled in 10 minutes was 4.98 ± 0.60 m (n=6) for littermate controls and 4.13 ± 0.49 m (n=6) for Munc18-1SA mice (p= 0.3). Both genotypes spent similar amounts of time in the center (WT: 9.3 ± 2.5%, n=6 and Munc18-1SA: 11.7 ± 3.8%, n=6; p= 0.6).

These findings indicate that Munc18-1SA mice are global knockin mice in which wildtype Munc18-1 has been replaced by Munc18-1SA, and expression levels of Munc18-1 are unchanged. These mice are therefore well suited to our primary goal, which was to determine the effect of Munc18-1 phosphorylation on synaptic transmission. The observation that in the absence of Cre the Munc18-1cSA mice are hypomorphs indicates that this mouse is not suitable for our secondary goal, which was to have a mouse that would allow us selectively replace Munc18-1 with Munc18-1SA in a Cre-dependent manner while leaving WT Munc18-1 at normal levels in other cells.

### Basal Properties of Synaptic Transmission

Prior to examining PTP, we characterized basal synaptic properties. We began by characterizing paired-pulse plasticity. At both CA3 to CA1 synapses and PF to PC synapses, the second of two closely spaced stimuli evokes a larger response than the first as a result of facilitation. Changes in the paired-pulse ratio (PPR) could arise either by directly contributing to facilitation, or indirectly by altering the initial probability of release (Zucker and Regehr, 2002). PKC phosphorylation of Munc18-1 has not been implicated in facilitation, which is thought to be mediated by Synaptotagmin 7 at the CA3 to CA1 synapse (Jackman et al., 2016). Alternatively, the most common means of altering PPR is by changing the initial probability of release (*p*): an increase in initial *p* indirectly decreases PPR and vice versa, in part because vesicle depletion is more prominent when initial *p* is high (Zucker and Regehr, 2002). We stimulated synaptic inputs with pairs of pulses separated by 20 to 600 ms. At the CA3 to CA1 synapse, for stimuli separated by 20 ms the PPR was 1.78 ± 0.06 in littermate controls (N= 4 animals, n=12 cells), and was moderately but statistically significantly reduced in Munc18-1SA mice (1.57 ± 0.06, N=4, n=13, **Fig. 2A, Table 2-1**). At the PF to PC synapse, the magnitude of facilitation was also statistically significantly reduced from 2.12 ± 0.07 (N=7, n=21) in littermate controls to 1.79 ± 0.07 (N=4, n=12) in Munc18-1SA mice (**Table 2-1**). These findings are consistent with a slight increase in the basal probability of release, or direct effects on paired-pulse plasticity at these synapses in Munc18-1SA mice.

**Figure 2.**
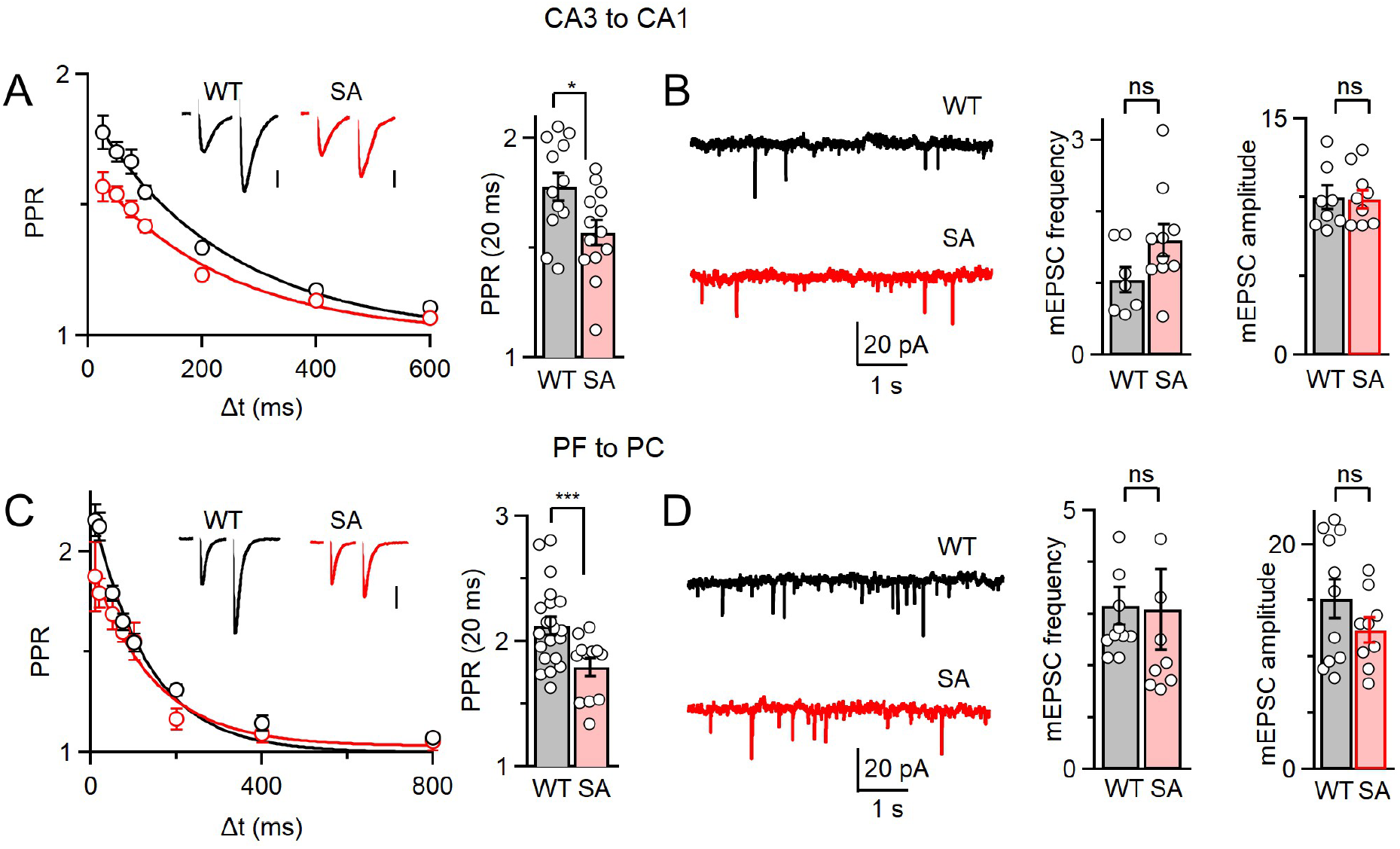
Basal synaptic properties in Munc18-1 SA-KI mice. A. Paired-pulse facilitation in wildtype and Munc18-1 SA KI mice at CA3 to CA1 synapses. Insets show representative synaptic currents evoked by pairs of stimuli separated by 20 ms. The decay of PPR was fit with a single exponential with a time constant of 240 ms for WT and 230 ms for Munc18-1 SA mice. Vertical scale bars are 100 pA. B. Example spontaneous mEPSCs in wildtype and Munc18-1 SA KI mice (left) and summaries of mEPSC frequencies and amplitudes (right). C,D. Same as A,B but for the cerebellar granule cell to Purkinje cell synapse. The decay of PPR was fit with a single exponential with a time constant of 129 ms for WT and 143 ms for Munc18-1 SA mice.

**Table 2-1.**
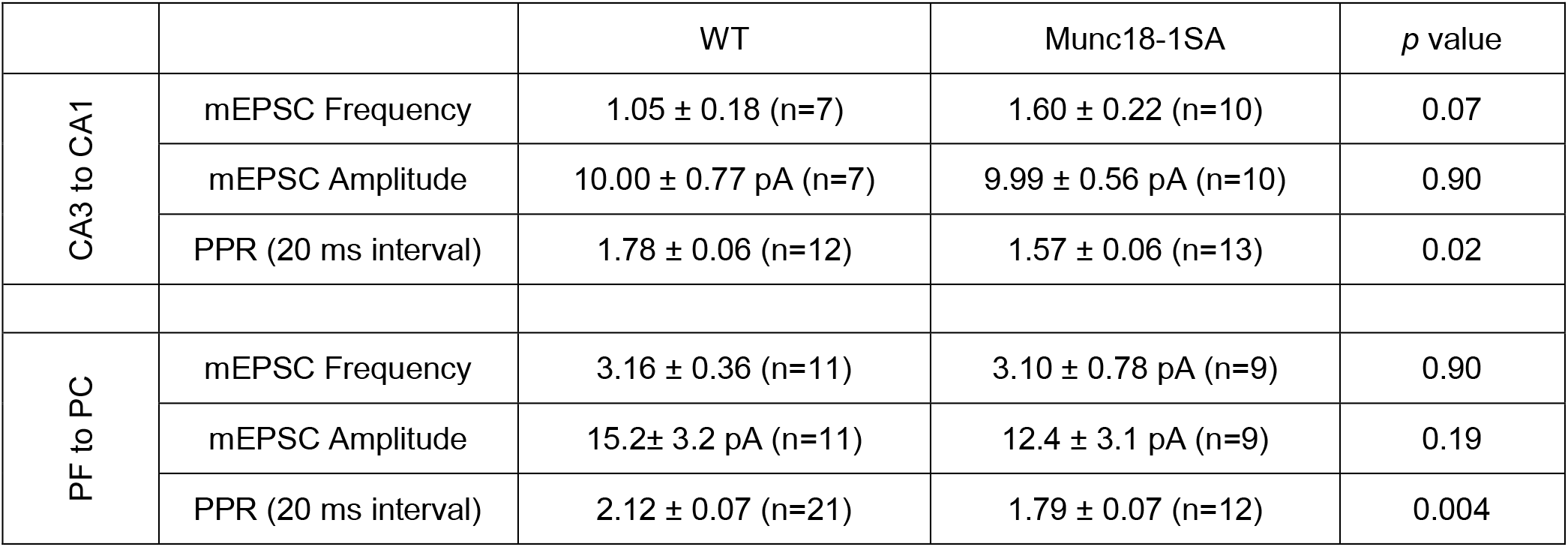
Data summary for Figure 2.

|  |  | WT | Munc18-1SA | *p* value |
| --- | --- | --- | --- | --- |
| CA3 to CA1 | mEPSC Frequency | 1.05 ± 0.18 (n=7) | 1.60 ± 0.22 (n=10) | 0.07 |
|  | mEPSC Amplitude | 10.00 ± 0.77 pA (n=7) | 9.99 ± 0.56 pA (n=10) | 0.90 |
|  | PPR (20 ms interval) | 1.78 ± 0.06 (n=12) | 1.57 ± 0.06 (n=13) | 0.02 |
| PF to PC | mEPSC Frequency | 3.16 ± 0.36 (n=11) | 3.10 ± 0.78 pA (n=9) | 0.90 |
|  | mEPSC Amplitude | 15.2 ± 3.2 pA (n=11) | 12.4 ± 3.1 pA (n=9) | 0.19 |
|  | PPR (20 ms interval) | 2.12 ± 0.07 (n=21) | 1.79 ± 0.07 (n=12) | 0.004 |

We also tested the possibility that the initial probability of release was altered in Munc18-1SA mice by determining the input-output curve at the CA3 to CA1 synapse (**Fig. 2-1**). We stimulated CA3 axons with a range of intensities and recording the resulting responses. The presynaptic volley provides a measure of the number of activated fibers, and the field excitatory postsynaptic potential (fEPSP) provides a measure of synaptic strength (Dingledine and Somjen, 1981). The ratio of the amplitude of the fEPSP/ the amplitude of the presynaptic volley was 1.88±0.29 (n=14) for WT mice and 2.46±0.20 (n=14) for Munc18-1SA mice, which are not significantly different (P=0.38, Wilcoxon rank sum test).

**Figure 2-1.**
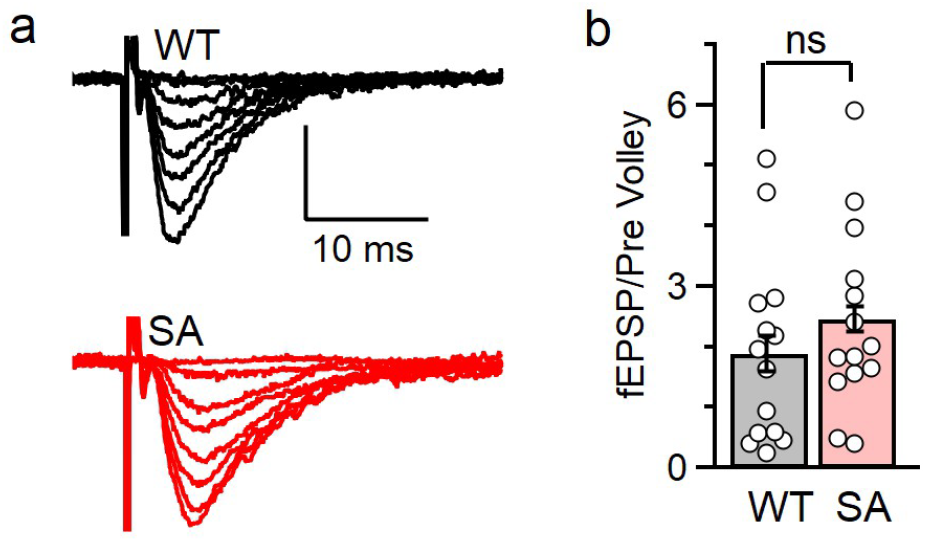
Input/output curves for the CA3 to CA1 synapse in wildtype and Munc18-1 SA mice. A. Representative field recordings of presynaptic volleys and fEPSPs for WT and Muun18SA mice evoked by a range of stimulus intensities. B. Summary of slopes of ratio of the field EPSP / presynaptic volley amplitudes. Bar graphs show the averages and SEM for WT (n=14, N=5) and Munc18SA (n=14, N=5) mice.

We also examined spontaneous neurotransmitter release of miniature excitatory postsynaptic currents (mEPSCs) in the presence of TTX. There was no statistically significant difference in either the amplitudes or the frequencies of mEPSCs for littermate controls and Munc18-1SA mice for CA3 to CA1 synapses (**Fig. 2A**) and for PF to PC synapses (**Fig. 2C**). Thus, the basal properties of spontaneous neurotransmitter release are not altered at these synapses in Munc18-1SA mice (**Table 2-1**).

### Phorbol esters-mediated enhancement

PKC can be activated by either calcium ions or by the second messenger diacylglycerol (DAG) produced by phospholipase C (PLC) (Zeng et al., 2012). The phorbol ester PDBu mimics DAG, and activates PKC by binding to the C1 domain of PKC. It has been proposed that PDBu enhances transmission by activating PKC and phosphorylating Munc18-1 in much the same way that elevations of presynaptic calcium can enhance transmission in PTP. At many synapses, phorbol esters both enhance transmission and occlude PTP (Korogod et al., 2007; Lou et al., 2005; Malenka et al., 1986; Saitoh et al., 2001; Shapira et al., 1987). This implicates PKC in both forms of enhancement, but interpretation of such occlusion experiments is complicated because PDBu can also activate proteins other than PKC that contain a C1 domain, such as Munc13 (Broeke et al., 2010; Brose and Rosenmund, 2002; De Jong et al., 2016; Hori et al., 1999; Junge et al., 2004; Lou et al., 2008; Rhee et al., 2002; Wierda et al., 2007).

Previous studies concluded that phorbol ester-mediated enhancement of mEPSC frequency was entirely dependent on PKC phosphorylation of Munc18-1 for cultured hippocampal cells (Wierda et al., 2007), but at the calyx of Held only half of the enhancement was mediated by this mechanism (Genç et al., 2014). These studies either expressed a PKC-insensitive Munc18-1 in a null background for cultured hippocampal cells, or conditionally eliminated Munc18-1 and virally expressed a PKC-insensitive Munc18-1 at the calyx of Held. Here we use Munc18-1SA mice to perform similar experiments at CA3 to CA1 synapses and PF to PC synapses. We found that at the CA3 to CA1 synapse PDBu produced large increases in the mEPSC frequency in both control (9.04 ±0.83-fold increase, N=2, n=10) and Munc18-1SA mice (12.82 ± 0.75-fold, N=2, n=10) (**Fig. 3A, Table 3-1**). There was no statistically significant difference in the magnitude of enhancement in littermate controls and Munc18-1SA-KI mice (p=0.30). We found that PDBu also increased mEPSC frequency at the PF to PC synapse in both littermate controls (2.53 ± 0.56-fold) and Munc18-1SA mice (3.88 ± 0.26, N=2, n=7) (**Fig. 3C**). This indicates that PKC phosphorylation of Munc18-1 does not account for enhancement by phorbol esters at the PF to PC synapse.

**Figure 3.**
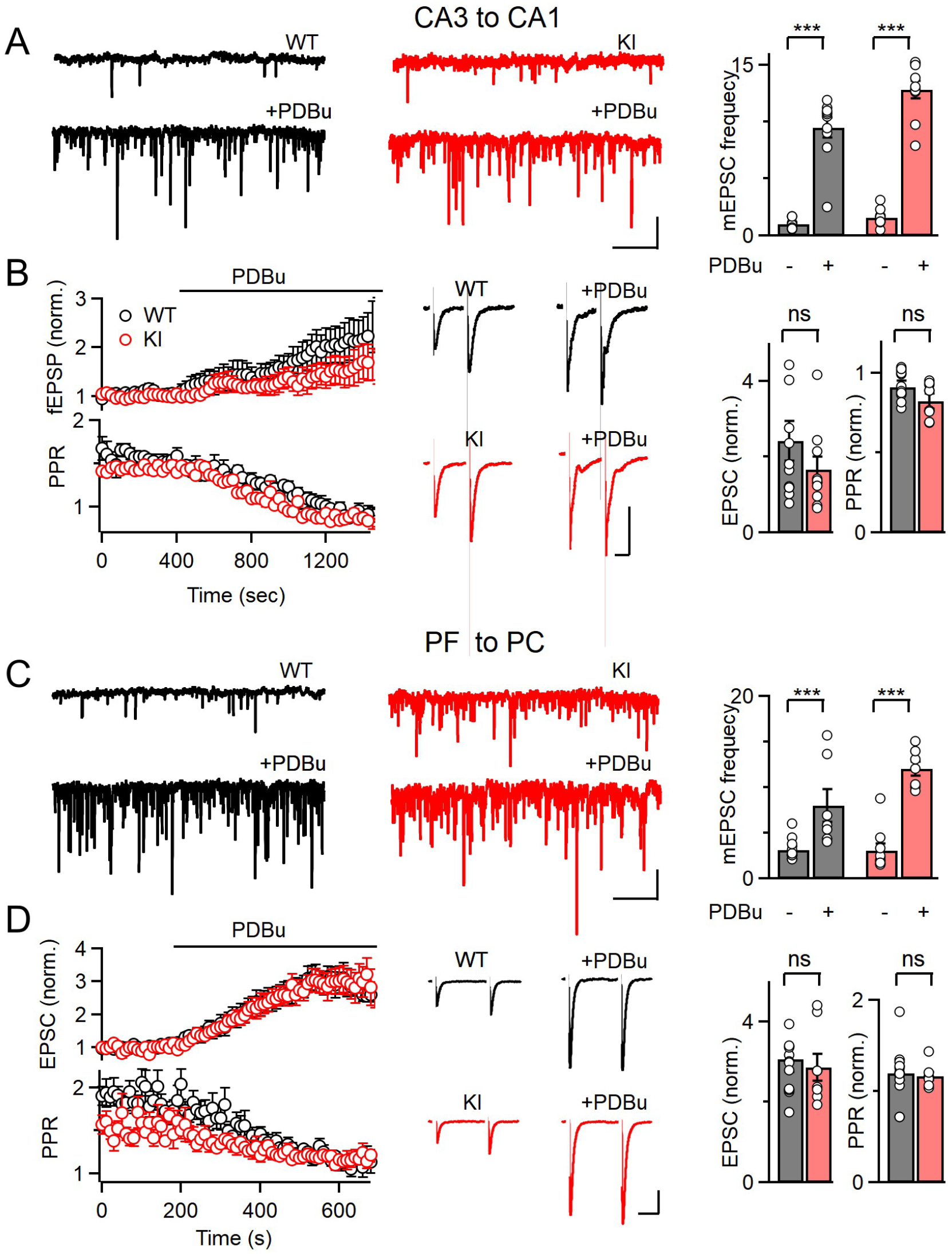
Phorbol esters enhance synaptic transmission in Munc18-1 SA-KI mice. A. The effect of the phorbol ester PDBu on mEPSCs recorded in CA1 pyramidal cells in the presence of TTX are shown for representative experiments before and during PDBu application (left and middle), and bare graphs summarize the results (right). B. The effects of phorbol ester PDBu on fEPSP amplitudes and paired pulse plasticity at CA3 to CA1 synapses for wildtype and Munc18-1 SA-KI mice. Summaries showing wash-ins (left), traces from representative experiments (middle), and bar graph summaries (right) are shown. C-D. Same as A, B but for Purkinje cells and the granule cell to PC synapse. In D EPSCs were measured in voltage clamp.

**Table 3-1.**
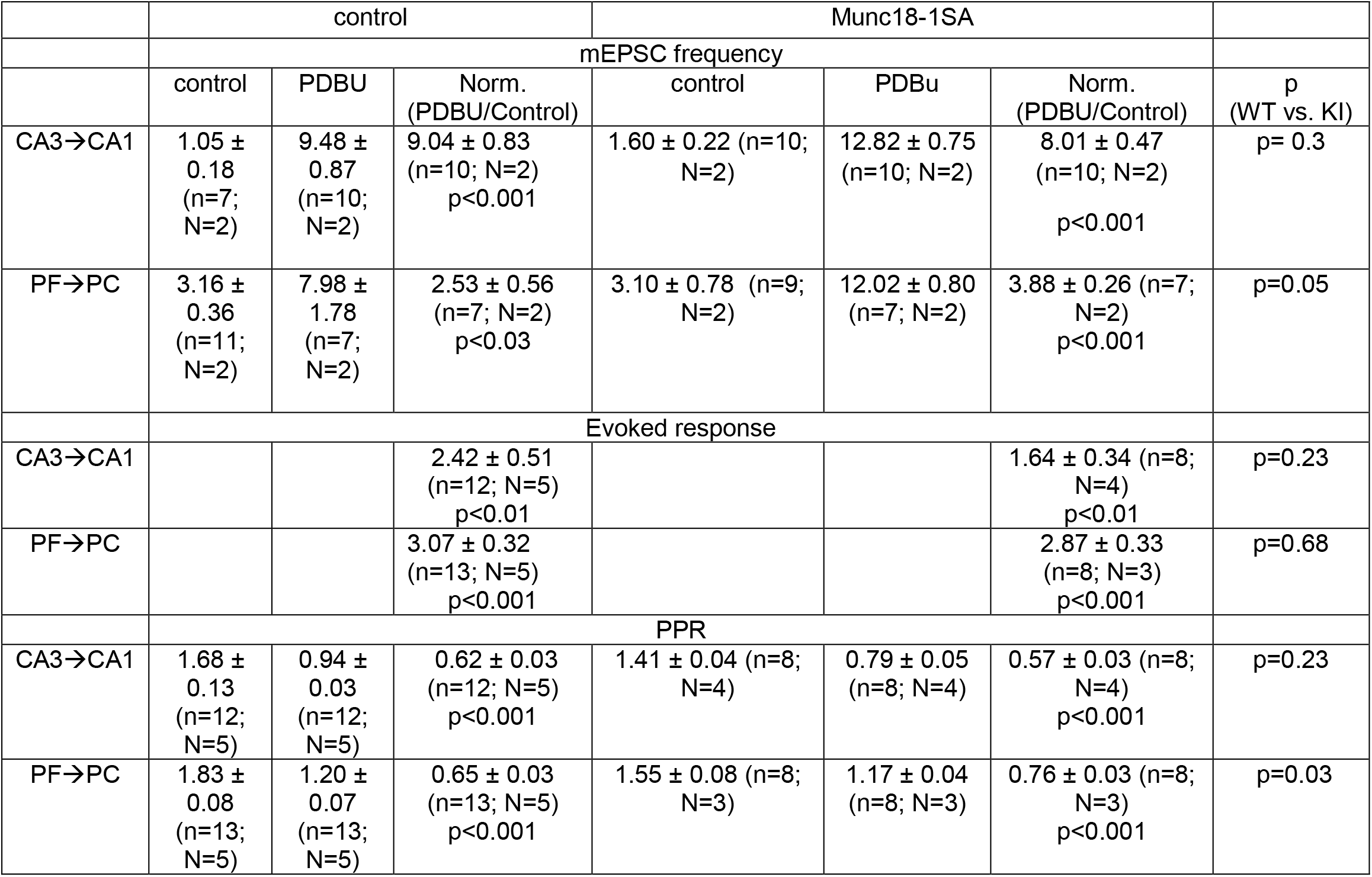
Data summary for figure 3: Summary of the effects of phorbol esters on spontaneous and evoked transmission at CA3 to CA1 synapses and PF to PC synapses.

|  | control |  |  | Munc18-1SA |  |  |  |
| --- | --- | --- | --- | --- | --- | --- | --- |
|  | mEPSC frequency |  |  |  |  |  |  |
|  | control | PDBU | Norm.<br>(PDBU/Control) | control | PDBu | Norm.<br>(PDBU/Control) | p<br>(WT vs. KI) |
| CA3→CA1 | 1.05 ± 0.18<br>(n=7; N=2) | 9.48 ± 0.87<br>(n=10; N=2) | 9.04 ± 0.83<br>(n=10; N=2)<br>p<0.001 | 1.60 ± 0.22 (n=10; N=2) | 12.82 ± 0.75<br>(n=10; N=2) | 8.01 ± 0.47<br>(n=10; N=2)<br>p<0.001 | p= 0.3 |
| PF→PC | 3.16 ± 0.36<br>(n=11; N=2) | 7.98 ± 1.78<br>(n=7; N=2) | 2.53 ± 0.56<br>(n=7; N=2)<br>p<0.03 | 3.10 ± 0.78 (n=9; N=2) | 12.02 ± 0.80<br>(n=7; N=2) | 3.88 ± 0.26 (n=7; N=2)<br>p<0.001 | p=0.05 |
|  | Evoked response |  |  |  |  |  |  |
| CA3→CA1 |  |  | 2.42 ± 0.51<br>(n=12; N=5)<br>p<0.01 |  |  | 1.64 ± 0.34 (n=8; N=4)<br>p<0.01 | p=0.23 |
| PF→PC |  |  | 3.07 ± 0.32<br>(n=13; N=5)<br>p<0.001 |  |  | 2.87 ± 0.33<br>(n=8; N=3)<br>p<0.001 | p=0.68 |
|  | PPR |  |  |  |  |  |  |
| CA3→CA1 | 1.68 ± 0.13<br>(n=12; N=5) | 0.94 ± 0.03<br>(n=12; N=5) | 0.62 ± 0.03<br>(n=12; N=5)<br>p<0.001 | 1.41 ± 0.04 (n=8; N=4) | 0.79 ± 0.05<br>(n=8; N=4) | 0.57 ± 0.03 (n=8; N=4)<br>p<0.001 | p=0.23 |
| PF→PC | 1.83 ± 0.08<br>(n=13; N=5) | 1.20 ± 0.07<br>(n=13; N=5) | 0.65 ± 0.03<br>(n=13; N=5)<br>p<0.001 | 1.55 ± 0.08 (n=8; N=3) | 1.17 ± 0.04<br>(n=8; N=3) | 0.76 ± 0.03 (n=8; N=3)<br>p<0.001 | p=0.03 |

At the CA3 to CA1 synapse, there was no statistical difference in the extent of PDBu enhancement of evoked EPSCs in control (2.42 ± 0.52 N=5, n=12) and Munc18-1SA mice (1.64 ± 0.33 N=4, n=10, p=0.23) (**Fig. 3B**). PDBu also decreased the PPR to a similar extent in control and Munc18-1SA mice (**Fig. 3B**). At the PF to PC synapse, PDBu enhanced transmission by 3.07± 0.32-fold (N=6, n=13) in wildtype animals and by 2.87± 0.33-fold (N=3, n=7) in Munc18-1SA mice. Our findings indicate that PKC phosphorylation of Munc18-1 plays either a modest role or no role in PDBu-dependent enhancement of spontaneous and evoked synaptic transmission at CA3 to CA1 and PF to PC synapses.

### PTP

We examined the role of Munc18-1 phosphorylation in PTP at the CA3 to CA1 synapse. At this synapse, a pharmacological study previously implicated PKC in PTP at this synapse (Brager et al., 2003), but we recently found that PTP is not dependent on calcium-dependent PKC isoforms (Wang et al., 2016). We found that 50 stimuli at 50 Hz produced synaptic enhancement (increased by 67± 9%, N=4, n=11) that decayed with a time constant of 9.9 ± 0.7 s (**Fig. 4A, C, Table 4-1**). PTP was accompanied by a decrease in PPR such that PPR 5 s after tetanic stimulation divided by the PPR prior to tetanic stimulation was 0.79 ± 0.02, and it recovered with a time constant (τ) of 11.1 s. This decrease in PPR is consistent with a transient elevation in release probability that has been shown to accompany PTP at some synapses (Zucker and Regehr, 2002). The amplitude of PTP was significantly different in Munc18-1SA animals (46 ± 5% enhancement, N=5, n=16, p=0.04), but PTP was significantly shorter lived (τ=7.6 ± 0.5 s, p=0.0095), and was also accompanied by a decrease in PPR (0.80±0.01, τ=9.1 s) (**Fig. 4B, C**). These results indicate that although there is a significant decrease in the amplitude and time course of PTP, most PTP at the CA3 to CA1 synapse is not reliant on PKC phosphorylation of Munc18-1.

**Figure 4.**
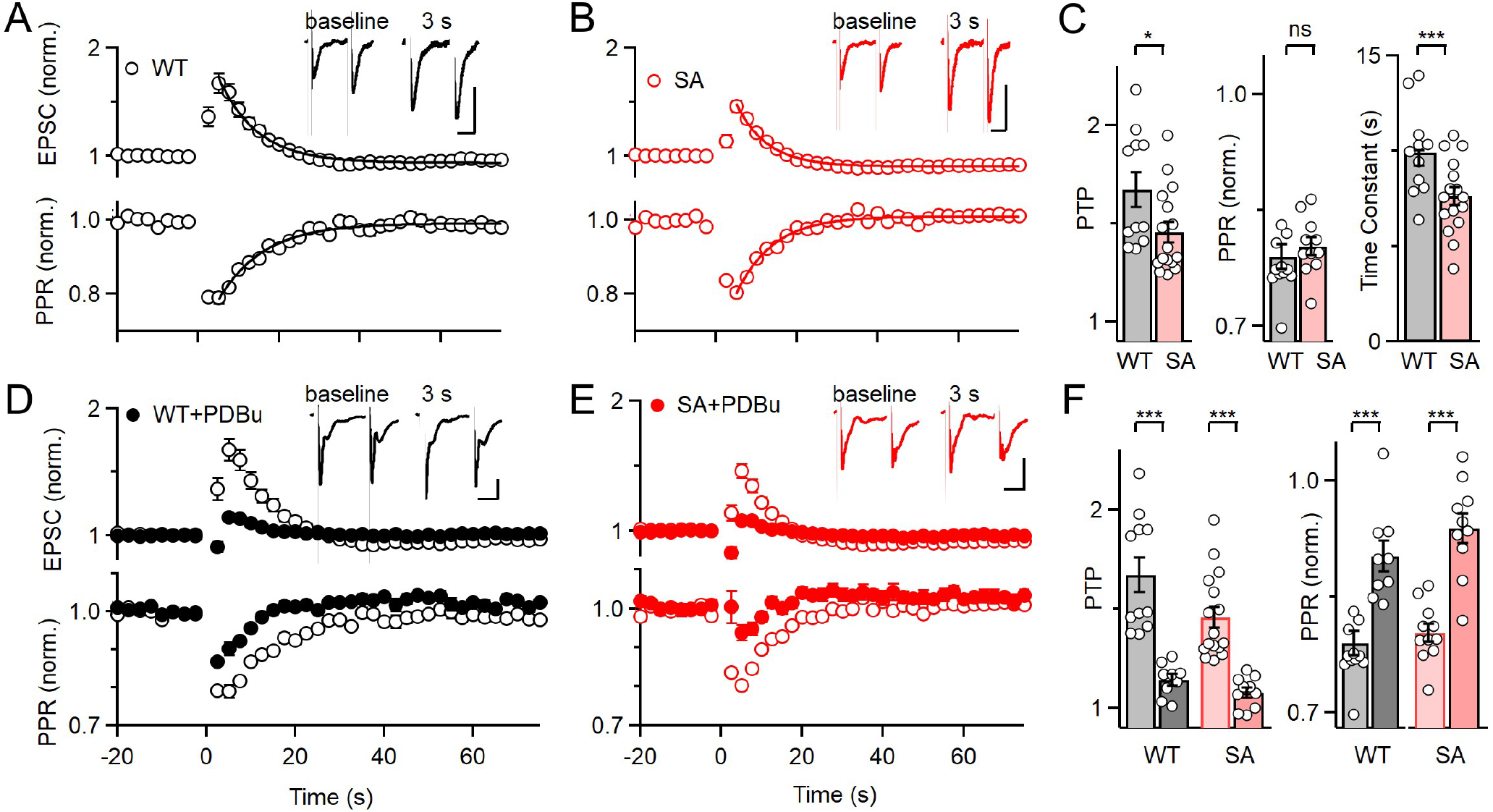
PTP is present at CA3 to CA1 synapses in Munc18-1 SA mice. A. PTP evoked by 50 stimuli at 50 Hz in WT mice with fEPSP amplitudes and paired-pulse plasticity summarized. Inset shows responses to pairs of stimuli prior to tetanic stimulation (*grey traces*) and 3 s. after (*black traces*). Scale bar: 0.2mV, 20ms B. Same as in A but for Munc18-1 SA mice (red), with the WT data replotted (black). The decays of PTP and changes in PPR were approximated with exponential fits (see text). C. Summary of individual experiments for A and B. D. PTP in wildtype mice measured in control conditions (open symbols) and in the presence of PDBu (solid symbols). E. As in D but for Munc18-1 SA mice. F. Summary of individual experiments for D and E.

**Table 4-1.**
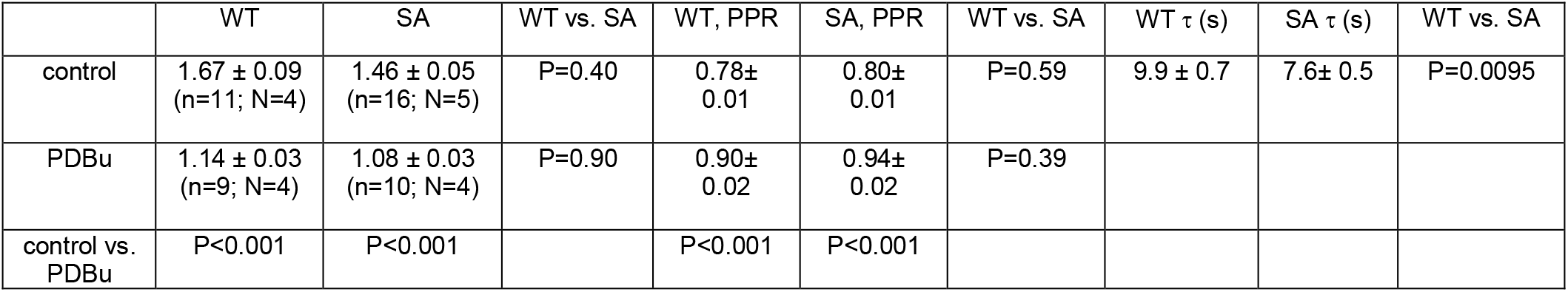
Data summary for Figure 4: Magnitude of PTP at CA3 to CA1 synapses data summary.

**Table 5-1.** Data summary for Figure 5: Magnitude of PTP at granule cell PF to PC synapse data summary (t=4.1s).

|  | WT | SA | WT (GF) | SA (GF) | TKO | TKO (GF) |
| --- | --- | --- | --- | --- | --- | --- |
| PTP | 1.717± 0.071<br>(n=19; N=8) | 2.146 ± 0.145<br>(n=18, N=8) | 1.089 ± 0.074<br>(n=17 N=8) | 1.265± 0.076<br>(n=17, N=7) | 1.677±<br>0.0689<br>(n=26, N=8) | 1.738± 0.166<br>(n=10, N=3) |

We also assessed the occlusion of PTP by phorbol esters at CA3 to CA1 synapses. The observation that phorbol esters occlude PTP could either arise from PKC activation and Munc18-1 phosphorylation (Genç et al., 2014; Wierda et al., 2007), or from an alternative mechanism such as PKC phosphorylation of another protein or phorbol ester activation of another protein (Brose and Rosenmund, 2002; de Jong and Fioravante, 2014; De Jong et al., 2016; Rhee et al., 2002). To determine if PDBu occludes PTP by phosphorylating Munc18-1, we compared the occlusion in littermate controls and Munc18-1SA mice (**Fig. 4D-F**). PTP was much smaller in the presence of PDBu for both wildtype mice (PDBu: 1.14 ± 0.03, n=9, N=4; vehicle: 1.67 ± 0.09, n=11, N=4) and Munc18-1SA mice (PDBu: 1.08 ± 0.03, n=10, N=4; vehicle: 1.46 ± 0.05, n=16; N=5). These findings indicate that PDBu occlusion of PTP at this synapse is not a result of phorbol esters activating PKC to phosphorylate Munc18-1.

We went on to examine PTP at the PF to PC synapse, where PKC has also been implicated in PTP. PTP was induced by a rather short stimulus train (10 stimuli at 50 Hz) to avoid producing presynaptic LTP (**Fig. 5A**). PTP was still present in Munc18-1SA animals with peak levels of PTP slightly larger than in wildtype animals (**Fig. 5B,D**). Thus, preventing PKC phosphorylation of Munc18-1 does not reduce PTP at the PF to PC synapse.

**Figure 5.**
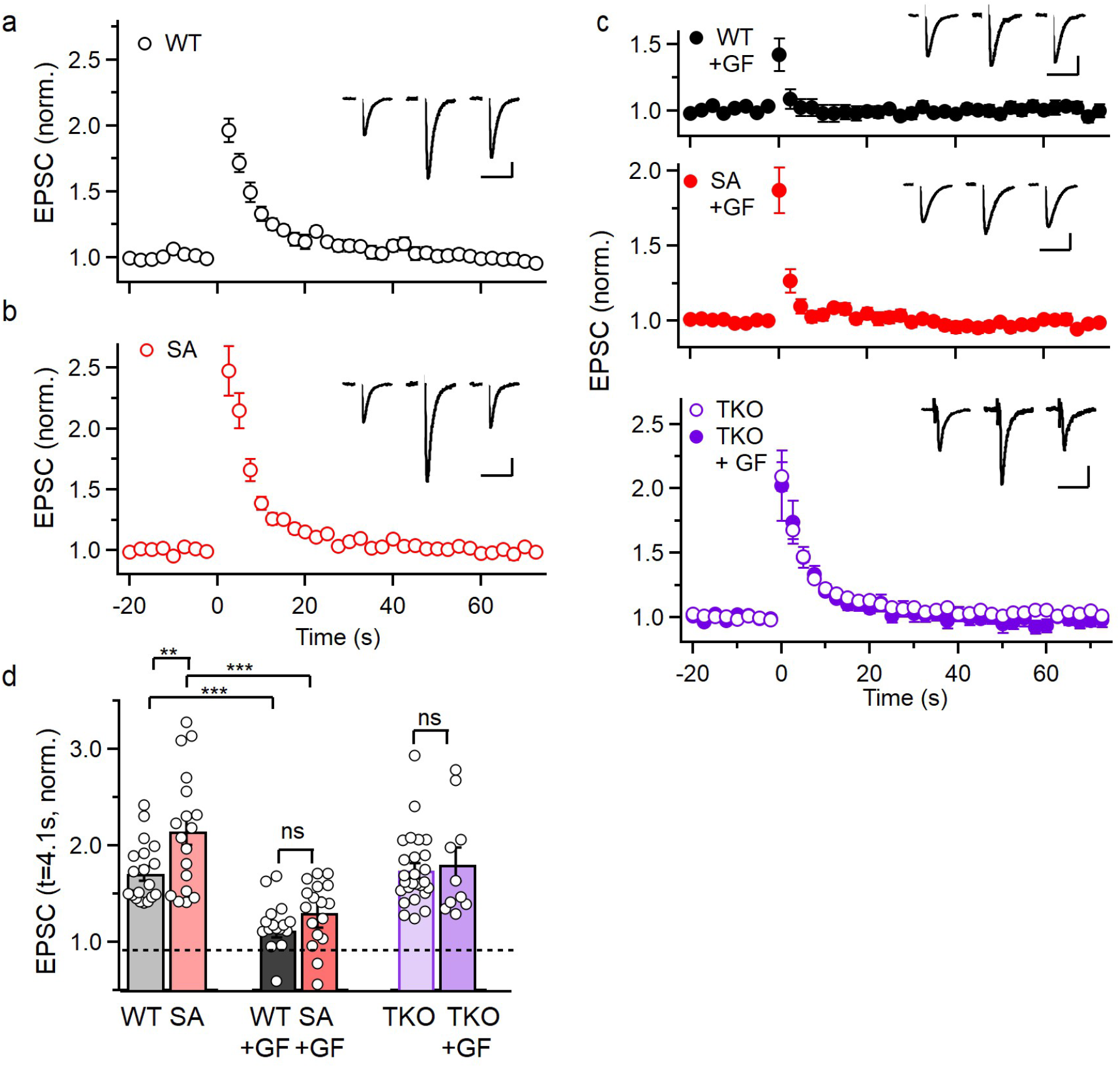
At PF to PC synapses most PKC-dependent PTP does not rely on Munc18-1. phosphorylation. PTP evoked by 10 stimuli at 50 Hz was examined at the granule cell to Purkinje cell synapse. a. Wildtype animals in control conditions (open black symbols). Inset shows example EPSCs recorded at −2.5 s, 4.1 s, and 60s. b. PTP in Munc18-1 SA mice. Scale bar: 200pA, 20ms c. The effect of 10 stimuli at 50 Hz on EPSC amplitude in the presence of the broad-spectrum PKC inhibitor GF (solid symbols) is shown for WT, SA and PKCαβγTKO mice. Insets show EPSCs recorded in the presence of GF. PTP induced in PKCαβγTKO mice in control conditions is included for comparison. d. Summary of effects of 10 stimuli at 50 Hz on EPSC amplitude for the indicated conditions. The second point after tetanic stimulation is plotted to reduce the effects of short-lived plasticity that persists in the presence of GF.

PTP at PF to PC synapses can also be mediated by a compensatory mechanism that is independent of PKC (Fioravante et al., 2012). This is illustrated by comparing the effect of the broad-spectrum PKC inhibitor GF109203X (GF) on PTP in wildtype animals and animals without calcium-dependent PKC isoforms. GF eliminates PTP in wildtype animals, which suggests that PKC is required for PTP at this synapse in wildtype animals. When calcium-dependent PKC isoforms are removed in PKCα, PKCβ, PKCγ triple knockout animals (PKCαβγ TKO animals), PTP is still present, but it is insensitive to GF (**Fig. 5C**,**D**), which is similar to PKCαβ DKO animals (Fioravante et al., 2012). These findings suggest that PTP is mediated by a PKC-dependent mechanism in wildtype animals, and by a compensatory PKC-independent independent mechanism in PKCαβγ TKO animals, (as in PKCαβ DKO mice (Fioravante et al., 2012)). It is not clear whether the PTP present in Munc18-1SA animals is mediated by PKC phosphorylation of a target other than Munc18-1, or is mediated by a PKC-independent compensatory mechanism as in the PKCαβγ TKO animals. GF can be used to distinguish between these possibilities: if PTP in Munc18-1SA animals is blocked by GF it is consistent with the former possibility, whereas if PTP in Munc18-1SA animals is insensitive to GF it is consistent with the latter possibility.

We confirmed that GF strongly attenuates PTP in WT mice, and found that it also strongly attenuated PTP in Munc18-1SA animals (**Fig. 5C**). These experiments also revealed that the first EPSC recorded after tetanic stimulation was still enhanced in both wildtype and SA mice in the presence of GF. For that reason, we compared the second EPSC (t=4.1 s), which provides a better measure of the longer lasting enhancement typical of augmentation and PTP (**Fig. 5D**). This suggests that in Munc18-1SA animals at PF to PC synapses almost all of the PTP relies on PKC. A comparison of PTP in blind and non-blind WT and Munc18-1SA animals in control conditions and in the presence of GF illustrates the differences in amplitudes and time courses of PTP. These results indicate that at PF to PC synapses, PTP is not reliant on PKC phosphorylation of Munc18-1, and suggests that PKC produces PTP at this synapse by phosphorylating a different target.

## Discussion

We assessed a mechanism of PTP in which calcium enters the presynaptic terminal, activates calcium-dependent PKC, which in turn phosphorylates Munc18-1 to enhance synaptic transmission. Our findings indicate that at CA3 to CA1 synapses and PF to PC synapses, phorbol ester-dependent enhancement and PTP are mediated primarily by mechanisms that do not involve PKC phosphorylation of Munc18-1.

### PKC phosphorylation of Munc18-1 and basal transmission

Our findings indicate that there are small effects on basal transmission at both CA3 to CA1 synapses and PF to PC synapses in Munc18-1SA mice, but they are difficult to interpret. The lack of alterations in spontaneous release rates and in mEPSC sizes suggest that the number of release sites is not altered and there are no postsynaptic changes in Munc18-1SA mice. The small decrease in paired-pulse plasticity in Munc18-1SA mice could reflect either an increase in the basal probability of release or a direct effect on paired-pulse plasticity. An increase in the initial probability of release could arise in Munc18-1SA mice if a preexisting pool of phosphorylated Munc18-1 lowers the initial probability of release in control animals, or if mutations present in Munc18-1SA alter the interaction of Munc18-1 with syntaxin or other presynaptic proteins independently of PKC phosphorylation to increase the probability of release. However, although the input/output values at the CA3 to CA1 synapse were not significantly different in Munc18-SA mice, the slope was steeper in Munc18-1SA mice (consistent with a higher initial p). This is reminiscent of a previous study that found that Munc18-1SA also produced a small decrease in paired-pulse plasticity for cultured hippocampal cells (Wierda et al., 2007), but without altering initial EPSC size, and therefore suggested that the initial probability of release was unchanged. Although the precise manner in which paired-pulse plasticity is altered in Munc18-1SA mice remains uncertain, it is clear that any effect of Munc18-1 phosphorylation on basal synaptic transmission is small.

### PKC phosphorylation of Munc18-1 and PTP

Our results establish that most PTP at the CA3 to CA1 synapse is not reliant on PKC phosphorylation of Munc18-1. PKC was initially implicated in PTP at the CA3 to CA1 synapse by a pharmacological study showing that phorbol esters enhance transmission and occlude PTP, and PKC inhibitors block PTP (Brager et al., 2003). A subsequent study confirmed these findings in wildtype animals, but found similar results in PKCαβγ TKO mice that lack calcium-dependent PKC isoforms (Wang et al., 2016). This indicates that calcium-dependent PKC isoforms are not the calcium sensors for PTP at this synapse, and left open two possibilities: (1) calcium-insensitive PKC isoforms mediate PTP or (2) PKCs are not involved in PTP at this synapse, and phorbol esters and PKC inhibitors influence by acting on proteins other than PKC. The most straightforward interpretation of the reduction of PTP in Munc18-1SA animals is that a small component of PTP relies on non-classical PKCs phosphorylating Munc18-It is also possible that in Munc18-1SA animals PTP could require non-classical PKCs phosphorylating targets other than Munc18-1, or could be mediated by a PKC-independent mechanism.

PTP at PF to PC synapses is also mediated primarily by mechanisms that do not rely on PKC phosphorylation of Munc18-1, but the role of PKC and Munc18-1 differs from that present at the CA3 to CA1 synapse. At PF to PC synapses, when calcium-dependent PKCs are eliminated, PTP is still present. The PKC inhibitor GF blocks PTP when calcium-dependent PKCs are present, but not when they are absent ((Fioravante et al., 2012), Figure 5c). This suggests that PKC is essential for PTP in wildtype animals, but a compensatory PKC-independent mechanism mediates PTP in PKCαβγ TKO mice. The compensatory mechanism that mediates PTP in the absence of calcium-dependent PKCs is not known, and there are many candidates, including the hundreds of phosphorylation sites on presynaptic proteins that are up- or down-regulated in a calcium-dependent manner (Kohansal-Nodehi et al., 2016). In Munc18-1SA mice, PTP is intact at the PF to PC synapse and no such compensation is apparent. The PTP that remains is mostly blocked by GF, suggesting that it is mediated primarily by a PKC-dependent mechanism. It appears that most PTP is mediated by PKC, but presumably as a result of phosphorylation of a protein other than Munc18-1. There are many candidates known to be phosphorylated by PKC, including SNAP25 (Shimazaki et al., 1996), synaptobrevin (Nielander et al., 1995), liprin-a3 (Emperador-Melero et al., 2020; Kohansal-Nodehi et al., 2016), and synaptotagmin 1 (De Jong et al., 2016). Further studies are required to determine whether PKC phosphorylation of any of these targets contributes to PTP.

The modest contribution of PKC phosphorylation of Munc18-1 to PTP we describe here for CA3 to CA1 and PF to PC synapses contrasts with previous studies at cultured hippocampal synapses (Wierda et al., 2007) and at the calyx of Held in brain slice (Genç et al., 2014). In hippocampal cultures, PTP induced by 20 or 200 stimuli at 40 Hz was present at WT synapses but was completely eliminated at Munc18-1SA (referred to as M18-1_PKCi_) synapses. At the calyx of Held, Munc18-1 was eliminated in conditional KO mice by expressing Cre in presynaptic cells, and using viruses to express either WT Munc18-1 or Munc18-1SA. WT rescue was enhanced by 120% compared to 50% for the SA rescue, and PTP decayed much more rapidly for the SA rescue such that the 60% reduction in initial PTP amplitude corresponded to an 84% reduction in the cumulative PTP. It was therefore concluded that PTP at the calyx of Held was a consequence of PKC phosphorylation of Munc18-1, with phosphorylation and enhancement persisting until phosphatases dephosphorylated Munc18-1. These results suggested that PKC phosphorylation of Munc18-1 accounted for all of the PTP for cultured hippocampal synapses and most of the PTP at the calyx of Held. Our results combined with these previous results indicate that PKC phosphorylation of Munc18-1 makes a variable contribution to PTP at different synapses, and that this mechanism is not a universal mechanism responsible for the bulk of PTP at most synapses.

### PKC phosphorylation of Munc18-1 and enhancement by phorbol esters

There are also significant differences in the contribution of Munc18-1 phosphorylation to enhancement by phorbol esters at different synapses. Here we find that in Munc18-1SA mice the phorbol ester enhancement of evoked synaptic responses was reduced by ∼50% at the CA3 to CA1 synapse (Figure 3A) and essentially unchanged at the PF to PC synapse (Figure 3B). This compares to complete elimination in cultured hippocampal synapses (Wierda et al., 2007), and a reduction by ∼50% at the calyx of Held (Genç et al., 2014) for a similar manipulation. The phorbol ester enhancement of mEPSC frequency was reduced by ∼10% at the CA3 to CA1 synapse (Figure 3C), increased at PF to PC synapses (Figure 3D), completely eliminated in cultured hippocampal synapses (Genç et al., 2014; Wierda et al., 2007), and reduced by approximately 50% at the calyx of Held (Genç et al., 2014). Thus, just as with PTP, the contribution of PKC phosphorylation of Munc18-1 to phorbol ester-mediated enhancement of transmission is highly synapse dependent.

### Challenges in determining the behavioral roles of PTP

It has been extremely difficult to determine the behavioral roles of PTP and other forms of short-term plasticity. This requires a means of selectively eliminating PTP, and ideally targeting specific types of synapses. Some of the challenges are apparent in considering PTP at hippocampal and cerebellar synapses. The implication of PKC in PTP suggested a target, but with PKC involved in many aspects of cell signaling more specificity was needed. The possibility that PTP is mediated by PKC phosphorylation of Munc18-1 suggested that replacing Munc18-1 with Munc18-1 SA could provide a means of selectively perturbing PTP. However, our studies have established that PTP is not eliminated at these synapses in Munc18-1 SA mice, and that these mice are not suitable for determining the behavioral role of PTP.

## Acknowledgements

This work was supported by grants from the National Institutes of Health (R01NS032405 and R35NS097284) to W.R., R01NS083898 to P.K., Goldenson Fellowship to C.C.W., a Boehringer Ingelheim Fonds PhD Fellowship, a B&C Privatstiftung Forschungsförderung, an Alice and Joseph Brooks Postdoctoral Fellowship to C.W.

